# Structure-guided design of a synthetic mimic of an EPCR-binding PfEMP1 protein

**DOI:** 10.1101/749432

**Authors:** Natalie M. Barber, Clinton K. Y. Lau, Louise Turner, Gareth Watson, Susan Thrane, John P.A. Lusingu, Thomas Lavstsen, Matthew K. Higgins

**Author notes:** Contributed equally (Natalie M. Barber, Clinton K. Y. Lau, Louise Turner).

## Abstract

Structure-guided vaccine design provides a route to elicit a focused immune response against the most functionally important regions of a pathogen surface. This can be achieved by identifying epitopes for neutralizing antibodies through structural methods and recapitulating these epitopes by grafting their core structural features onto smaller scaffolds. In this study, we have conducted a modified version of this protocol. We focused on the PfEMP1 protein family found on the surfaces of erythrocytes infected with *Plasmodium falciparum*. A subset of PfEMP1 proteins bind to endothelial protein C receptor (EPCR), and their expression correlates with development of the symptoms of severe malaria. Structural studies revealed the PfEMP1 to present a helix-kinked-helix motif which forms the core of the EPCR binding site. Using Rosetta-based design we successfully grafted this motif onto a three-helical bundle scaffold. We show that this synthetic binder interacts with EPCR with nanomolar affinity and adopts the expected structure. We also assessed its ability to bind to antibodies found in immunized animals and in humans from malaria endemic regions. Finally, we tested its capacity to effectively elicit antibodies that prevent EPCR binding and analysed the degree of cross-reactivity of these antibodies across a diverse repertoire of EPCR-binding PfEMP1. This provides a case study of immunogen design, assessing the effect of designing a focused immunogen that contains the core features of a ligand binding site, rather than those of a neutralizing antibody epitope.

## Introduction

Parasites must often expose proteins on their surfaces to mediate interactions with molecules of their mammalian host, allowing host cell invasion, nutrient uptake and modulation of host immunity. The evolutionary driving forces which shape these parasite surface proteins are multiple, with the need to retain the capacity to interact with an unchanging mammalian binding partner balanced against the pressure to resist detection and clearance by the acquired immune system of the host. A common solution is the evolution of a family of proteins, which are deployed one at a time by the parasite though antigenic variation (1–3). This allows a population survival strategy in which parasites which express a protein that is recognized by host immunoglobulin are cleared, while those that express a functionally equivalent, but unrecognized variant, survive. Such parasite surface protein families prove a major challenge to vaccine development, with their varying nature hampering efforts to design an immunogen which elicits a protective immune response across a family.

A classic example of a surface protein family is the PfEMP1 proteins of *Plasmodium falciparum* (1, 4). These are found on the surfaces of parasite-infected erythrocytes and interact with various human endothelial receptors. This causes infected erythrocytes to adhere to the surfaces of blood vessels and tissues, preventing their destruction by splenic clearance (5). Pathology in severe and placental malaria is associated with these adhesive properties, with endothelial binding occluding blood flow and inducing inflammatory responses (6). While there are many PfEMP1, and different family members bind to different human endothelial receptors (1, 4), a growing body of evidence suggests that severe and cerebral malaria is associated with expression of a subset of PfEMP1 that bind to endothelial protein C receptor (EPCR) (7–15). PfEMP1 prevents EPCR from binding to its natural ligand, protein C (7, 8), thereby preventing PAR1-mediated endothelial signaling (16), most likely resulting in inflammation and endothelial dysfunction. Indeed, the EPCR-binding PfEMP1 are targeted by antibodies which are found in adults from malaria endemic regions (8), are acquired early in life (17) and whose presence correlates with the individual having experienced a case of severe malaria (18). However, immunization of rodents with single CIDRα1 domains does not generate antibody responses against the full repertoire of EPCR-binding domains (19). This raises the question of whether it is possible to design vaccine immunogens which induce broadly inhibitory antibody responses that can target the sequence diverse set of EPCR-binding PfEMP1.

The ectodomains of PfEMP1 proteins are formed from an array of two domain types, the DBL and CIDR domains (3), which have been grouped into a variety of subclasses (20). PfEMP1 are generally modular, with single domains containing the capacity to bind to individual receptors (8, 9, 21). The majority of CIDRα1 domain subclasses bind to EPCR (7, 8). Structural studies of these domains in complex with EPCR, have revealed an interaction interface consisting of a hydrophobic core surrounded by a surface which mediates hydrogen bonds (8). At its centre, lies a helix-kinked-helix structural motif, which contains seven of the nine EPCR-interacting residues. At the kink lies a phenylalanine residue, F656 in HB3var03, which is central to the binding site and whose mutation leads to a 100-fold increase in the dissociation-rate of the complex (8).

Analysis of the sequences of 885 CIDRα1 domains reveals that the interacting residues are not conserved in sequence, but nevertheless maintain conserved chemical characteristics, with retention of the hydrophobic nature of the protrusion and the surrounding hydrophilic surface (8). This raises the question of whether it is possible to design a protein which mimics this surface, and if such a protein can specifically elicit antibodies which block EPCR binding. If so, such induced antibodies would be functionally valuable in preventing the modulation of EPCR-mediated signaling implicated in the development of severe symptoms. In addition, by targeting the most conserved part of the CIDRα1 domain surface, these antibodies would have the greatest likelihood of being cross-reactive across the EPCR-binding PfEMP1 family.

A recent advance in structure-guided immunogen design is an approach in which structures are determined for pathogen surface proteins in complex with antibodies with protective or broadly-neutralising properties, followed by the grafting of the epitopes of these antibodies onto smaller scaffolds (22, 23). In a number of cases, this has allowed the development of smaller immunogens which can specifically elicit the production of antibodies with desirable properties (24–27). The conserved chemistry and shape of the EPCR-binding site of the PfEMP1 (8) encouraged us to make a similar attempt. In this case, however, the absence of structural insight into the epitopes of inhibitory antibodies led us to trial a variation of this usual approach, in which we designed a smaller protein which mimics the features of the EPCR-binding site, while removing other potential epitopes found in CIDRα1 domains. In this we aimed to design a molecule which could be used as a tool, to assess the role of antibodies which target the core of the EPCR binding site. We also aimed to test this molecule as a vaccine immunogen to attempt to specifically elicit antibodies against the EPCR binding site and to assess their degree of cross-reactivity across the EPCR-binding PfEMP1.

## Results

### Iterative design of a novel EPCR-binding protein

Our design strategy was informed by structural studies of complexes of CIDRα1 domains bound to EPCR (8). Nine amino acids from CIDRα1 domains directly contact EPCR, with seven of these present in a motif consisting of a helix followed by a helix with a kink. We reasoned that we could graft this helix-kinked-helix motif onto a scaffold protein to generate an EPCR-binding protein containing the majority of the functional determinants of the CIDRα1 domain. To confirm that the two EPCR-contacting residues which would be missing from this design, D576A and K642A, were not essential for EPCR binding, we prepared mutated versions of the HB3var03 CIDRα1 domain in which they were replaced by alanine. The double mutant of these two residues showed a reduction in affinity, from 0.4nM to 220nM (Figure S1). As this nanomolar affinity was in line with the range of affinities of CIDRα1 domains for EPCR measured previously (8), we continued with the design process.

To select a suitable scaffold on which we could recapitulate this helix-kinked-helix motif, we searched known protein folds, using PDBeFold and DALI (28). However, this identified no template with a matching surface exposed structural motif. However, a three-helical bundle scaffold (PDB code 3LHP, chain S) used previously for immunogen grafting (29) contained α-helices whose path matched those of the two longer helices of the helix-kinked-helix motif, with the two helical portions overlaying with a RMSD of 2.5Å. We therefore grafted the helix-kinked-helix motif onto this scaffold and used a Rosetta-based strategy (25, 30) to redesign the resultant molecule to obtain the appropriate fold (Figure 1A).

**Figure 1:**
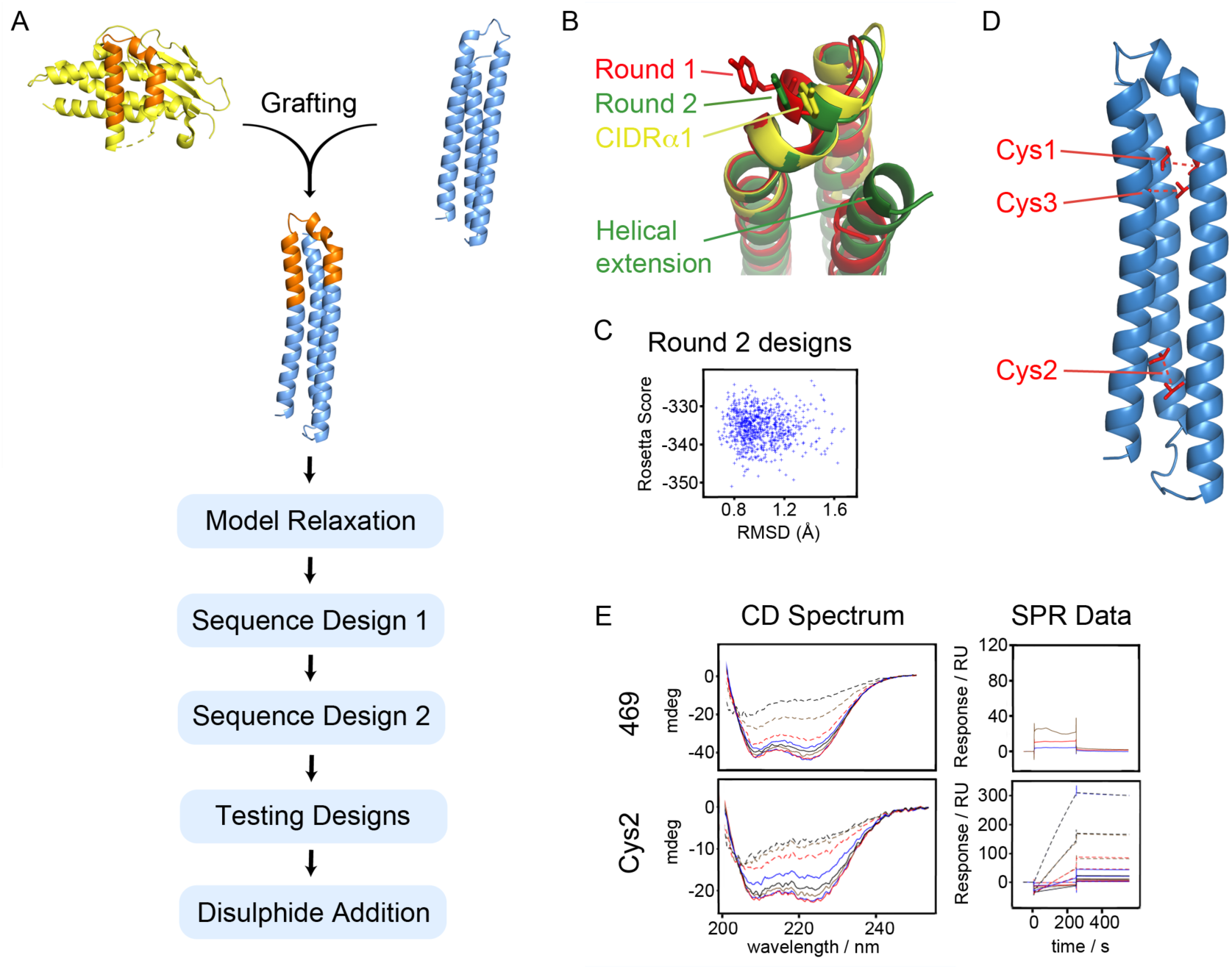
Design of a synthetic EPCR binder. **A.** A schematic showing the design process. The EPCR binding surface (orange) of the HB3var03 CIDRα1 domain (yellow) was grafted onto a three-helical bundle scaffold (blue), followed by a Rosetta-based design strategy. **B.** Illustration of the outcome of design rounds 1 (red) and 2 (green) illustrating the importance of increasing the length of the C-terminal helix in generating a design that more closely mimics the CIDRα1 domain (yellow). **C.** Analysis of output models from round 2, showing the Rosetta score and the root mean square deviation (RMSD) to the original epitope. **D.** A model of the designed binder, indicating positions of residues mutated to cysteine to introduce stabilizing disulphide bonds. **E.** Circular dichroism spectra and surface plasmon resonance analysis of EPCR-binding of the best design from round 2 (469) and the best disulphide-stabilised design (Cys2).

The starting model for the designed immunogen contained residues N648 to K678 from the HB3var03 CIDRα1.4 domain inserted in between residues Gly70 and Asp103 of the three-helical bundle (Figure 1A). This hybrid was assembled *in silico* and then computationally refolded in Rosetta by allowing other features of the immunogen to fold around a rigidly held epitope region (residues N648 to K678). This procedure was supplemented by Cα-Cα restraints derived from the original helical bundle to drive the model towards the target conformation. After generation of each model, restraints were removed, the epitope was unlocked, and the structure was allowed to relax to a local energy nadir. A plot of Rosetta score (a measure of the intramolecular interaction strength) against standard deviation to the starting model, generated two clusters, one with RMSD values of 0.5 – 2.5Å to the starting model and one at significantly higher RMSD, with a helix displaced. The model with the lowest RMSD was chosen for subsequent design.

We next used the fold-from-loops procedure to adjust the initial design to improve its folding. Each residue was classed as a ‘surface’, ‘boundary’ or ‘interior’ residue and was allowed to change to other amino acids within the same class. We prevented variation of eight residues which lie on the surface of the helix-kinked-helix in the CIDRα1 domain (corresponding to N648, D649, D652, S653, F655, F656, Q657 and Y660 from HB3var03 CIDRα1.4) and three residues which were thought likely to be important to maintain the conformation of the kinked helix (F651, V658 and W669). Four rounds of sequence design involved mutation followed by sequence relaxation in Rosetta and determination of Rosetta score. Early rounds were constrained by Cα-Cα restraints derived from the desired conformation of the helix-kinked-helix, but these constraints were removed in the final stage, allowing the models to adopt a minimum energy conformation. These models were assessed based on their Rosetta score and their root-mean-square deviation to the helix-kinked-helix motif. Assessment of the outputs from the first round of design shows deviations in the kinked helix from the original (Figure 1B). This was resolved by increasing the length of the third helix of the helical bundle, which lies behind the helix-kinked-helix motif to provide additional support. Performing the same process of sequence design using this longer scaffold generated models which more closely match the starting design (Figure 1B, C).

The design process generated a set of output models with as little as 35% pairwise sequence identity (Figure S2). To select which models to test experimentally, we selected the 100 best based on root-mean-square deviation to the epitope. These were classified by evolutionary trace analysis into eight groups, simply based on sequence, and a member of each of the eight groups was selected to give a low root-mean-square deviation to the epitope and a high packstat filter score (Figure 1C), indicating a well packed structure. These eight designs had pairwise sequence similarities of 35-60% (Figure S2).

Genes corresponding to these eight designs were ordered, with codon optimization for expression in *E. coli* and the inclusion of an N-terminal histidine tag (Table S1). Seven of these designs could be expressed in a soluble form and were purified. In each case, purification by size exclusion chromatography revealed a single monodispersed species. Circular dichroism spectroscopy revealed the seven designs to have a predominantly α-helical profile (Figure S3). Thermal melts, analysed by CD, revealed the temperatures at which α-helicity was lost and showed major variation in stability across the seven designs, with, for example, 398 retaining more than 90% of helical character at 90°C and 496 melting at around 70°C. In contrast, others, such as 555 shows a broad, non-cooperative transition towards loss of helicity, starting at 20°C (Figure S3). Next, to assess if the designs had adopted the correct fold, we assessed their ability to bind to EPCR as measured by surface plasmon resonance. However, while the parent HB3var03 CIDRα1.4 domain binds to EPCR with an affinity of 0.36nM, no binding was seen for any of the seven designs at 500nM, suggesting that the EPCR binding surface had not been correctly mimicked (Figure S3).

We reasoned that the inability of the seven designs to bind to EPCR might suggest a lack of structural stability. We therefore further stabilized the designs by the addition of disulphide bonds (Figure 1D). We selected three sites in design 469 at which residues were predicted to be found at an appropriate distance to allow disulphide bond formation. These residues were replaced with cysteine to form three mutants, cys1, cys2, and cys3, each of which had a single disulphide bond. These were tested as above (Figure S4). The cys2 variant was most effective, retaining α-helicity at up to 70°C in circular dichroism measurements and interacting with EPCR with a slow off rate, reminiscent of the CIDRα1-EPCR interaction (Figure 1E, Figure 2A-C). As determined by surface plasmon resonance, the cys2 variant bound to EPCR with an affinity of 26nM (Figure 2A), which compares with 0.4nM for HB3var03 CIDRα1.4 and 220nM for the D576A K642A mutant of this domain (Figure S1). We therefore proceeded with this design for structural and functional testing.

**Figure 2:**
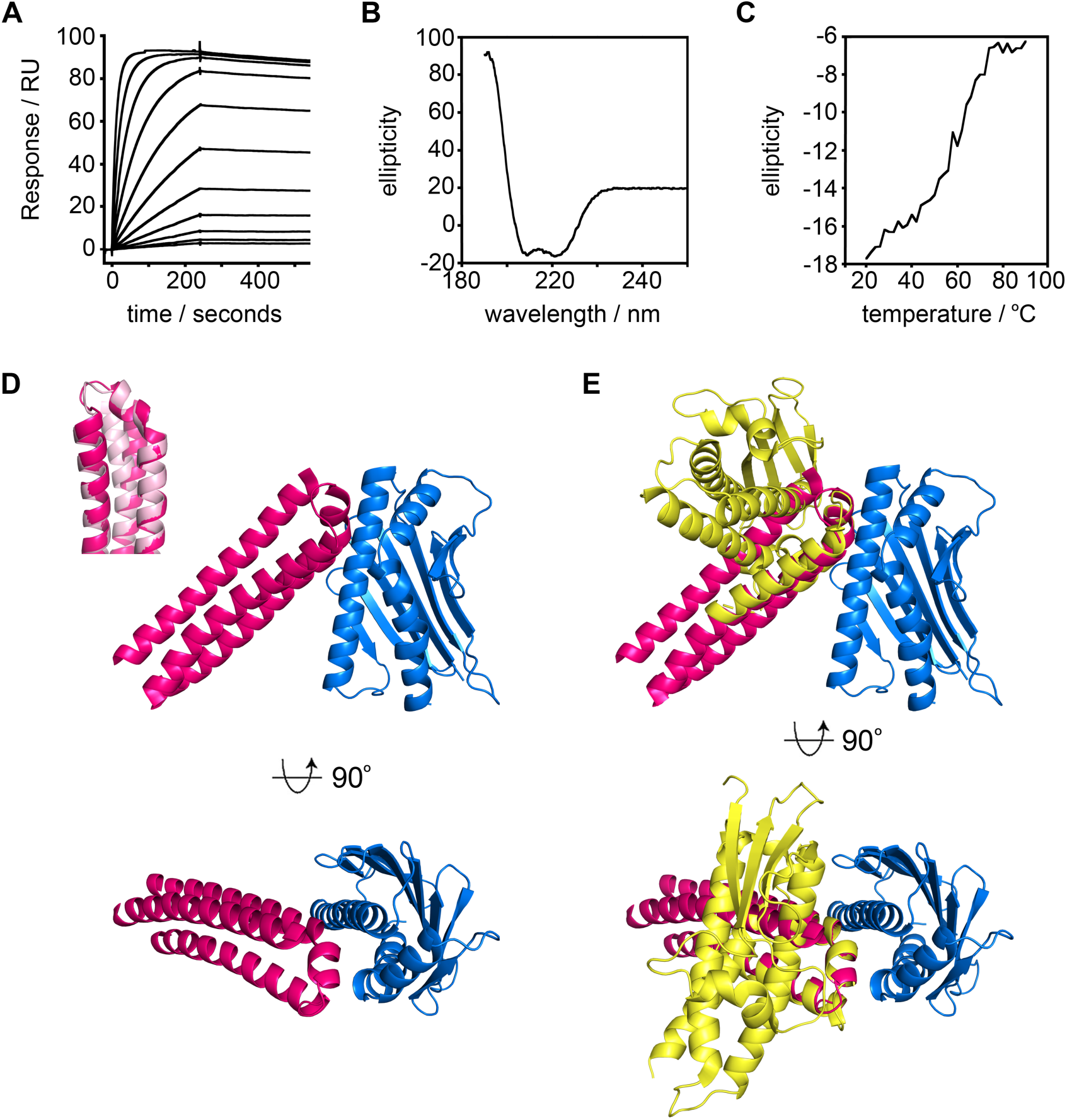
Structural and biophysical characterization of the synthetic binder. **A.** Surface plasmon resonance analysis of the binding of the synthetic binder to EPCR. The data shows a 2-fold dilution series starting from 16μM. **B.** Circular dichroism analysis of the synthetic binder. **C.** Thermal stability of the synthetic binder, determined by the circular dichroism signal at 222nm wavelength at different temperatures. **D.** Structure of EPCR (blue) bound to the synthetic binder (pink). The inset shows an overlay of the structure of the synthetic binder (pink) overlaid with the original design (light pink). **E.** An overlay of the structures of EPCR (blue) in complexes bound the synthetic binder (pink) and the HB2var03 CIDRα1 domain (yellow), with EPCR molecules overlaid.

### Structure of the synthetic EPCR binder in complex with EPCR

We next determined the crystal structure of the synthetic binder in complex with EPCR (Figure 2, Figure S5, Table S2). We cleaved purified synthetic binder and EPCR with TEV protease to remove tags, and deglycosylated EPCR. These were combined, and the complex purified by size-exclusion chromatography before being subjected to crystallization trials. Crystals formed and a complete data set was collected to 3.11Å resolution. Molecular replacement, using the EPCR structure (8) as a search model, identified two copies of EPCR in the asymmetric unit of the crystal. Surprisingly, a cycle of refinement and model building revealed the presence of a single copy of the helical bundle, with the asymmetric unit containing a complex of EPCR bound to a helical bundle, together with a second, un-liganded, copy of EPCR (Figure S5).

The structure of the helical bundle was compared with the model that emerged from the design process (Figure 2D). A structural alignment revealed the EPCR-binding site to adopt an extremely similar conformation to that predicted, with the residues of the helix-kinked-helix motif (N72-W93) overlaying with the design with a root-mean-square deviation of 1.19 Å. The structure also revealed the helical bundle to bind to EPCR with the same binding mode as the HB3var03 CIDRα1 domain (Figure 2E). Indeed, aligning these two complexes on the EPCR molecule, showed the residues of the helix-kinked-helix (N72 to W93) to overlay with the corresponding residues of the HB3var03 CIDRα1.4 domain with a root-mean-square deviation of 0.77Å. Therefore, the designed synthetic binder contains the core of the EPCR-binding site of the CIDRα1 domains and binds in the predicted manner to EPCR.

### Assessment of antibody binding to the synthetic binder

We next asked whether our synthetic binder interacts with either antibodies present in rats immunized with CIDRα1 domains, or with human antibodies from adult volunteers from malaria endemic regions of Tanzania. This would allow us to determine whether the helix-kinked-helix is the target of such antibodies and if these are inhibitory of EPCR binding.

Immunization of rats with the HB3var03 CIDRα1.4 domain induces the production of antibodies which prevent the cognate CIDRα1.4 domain from binding to EPCR (19) (Figure 3A). We affinity-purified antibodies from this serum on columns coupled with either HB3var03 CIDRα1.4 or our synthetic binder. Affinity purification on the CIDRα1 domain resulted in a flow-through that contained only ∼20% of the inhibitory activity found in the original sera, suggesting that the majority of inhibitory antibodies were depleted by binding to CIDRα1 domain. In contrast, passage through a column containing the synthetic binder did not reduce the capacity of the flow-through to inhibit EPCR binding by the CIDRα1 domain, suggesting that the majority of inhibitory antibodies in this serum do not bind to the synthetic binder. Nevertheless, the antibodies eluted from the synthetic binder did show some inhibitory capacity causing 40% inhibition of CIDRα1 domain binding to EPCR at a 50% dilution (Figure 3A). This could be compared with nearly complete inhibition of EPCR binding at a 3.1% dilution of antibodies purified on the CIDRα1 domain. Therefore, despite the synthetic binder adopting the correct confirmation, it is not recognized by the majority of CIDRα1 reactive antibodies that block EPCR binding in these sera.

**Figure 3:**
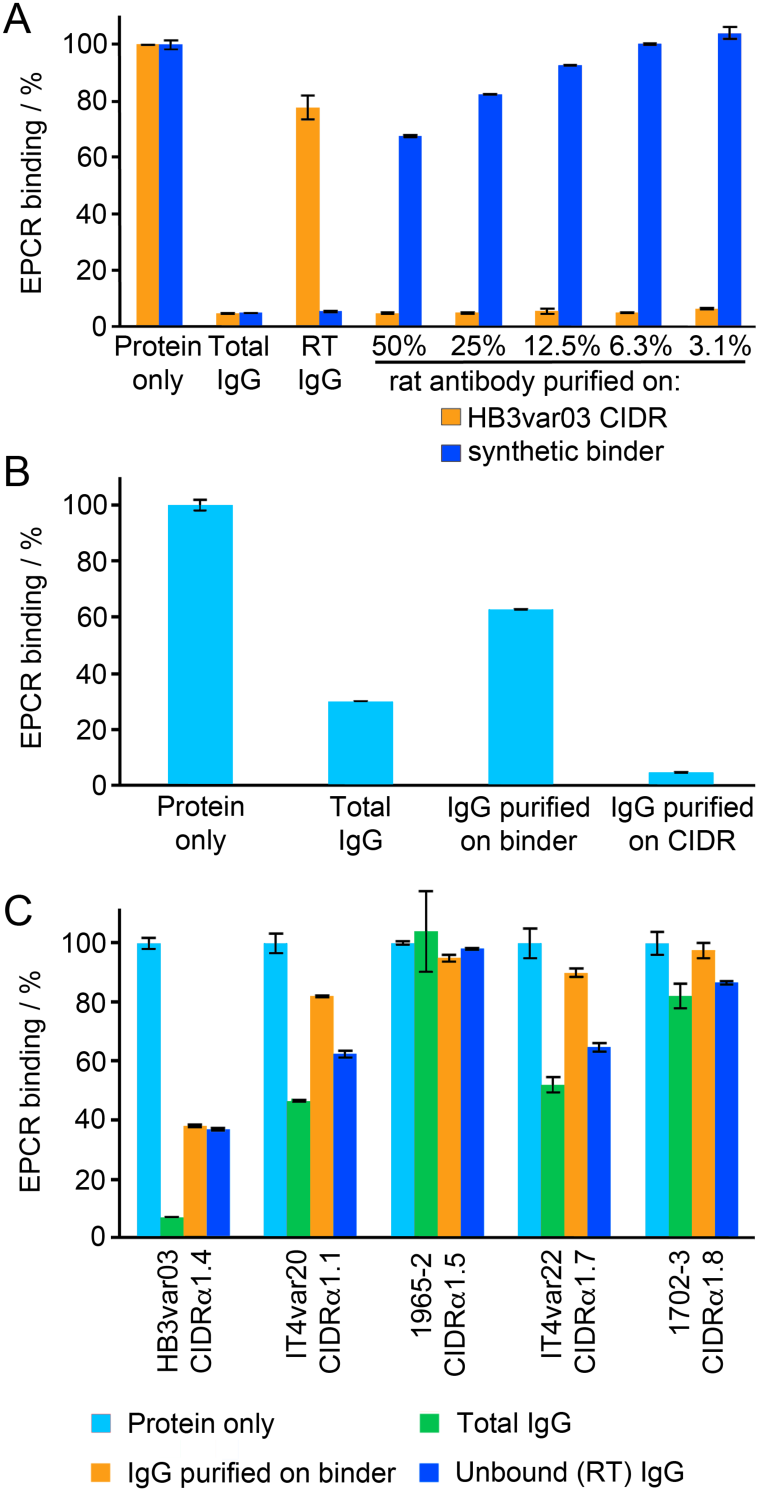
Targeting of the synthetic binder by antibodies from immunized rats or from humans from malaria endemic regions. **A.** Antibodies from rats immunized with the HB3var03 CIDRα1.4 domain were affinity purified using either HB3var03 CIDRα1.4 domain or the synthetic binder. These were assessed for their ability to prevent CIDRα1.4 domain from binding to EPCR. Protein alone indicates binding of EPCR in the absence of antibody. Total IgG shows EPCR binding in the presence of 0.55mg/ml total IgG. RT IgG shows EPCR binding in the presence of antibodies that did not bind to the affinity column. The remaining columns show EPCR binding in the presence of affinity purified antibodies at different dilutions. All data are expressed as a percentage of the binding in the absence of antibody. **B.** Prevention of CIDRα1 binding to EPCR by purified rat antibodies was quantified at 25µg/ml. Binding in the presence of Total IgG or of IgG affinity purified on the synthetic binder or the HB3var03 CIDRα1.4 domain was expressed as fraction of the binding in the absence of antibody (protein alone). **C.** Sera taken from fifteen individuals from malaria endemic regions of Tanzania were pooled and tested for their ability to prevent five different CIDRα1 domains from binding to EPCR. The total antibody pool was tested, as were antibodies purified on a column coupled with the synthetic protein and antibodies that did not bind to this column (RT IgG). In each case, this is expressed as a percentage of the binding in the absence of any antibody (protein alone).

We next quantified this effect by assessing the inhibitory capacity of the affinity purified antibodies at fixed concentrations of 25µg/ml (Figure 3B). Total purified IgG caused a ∼70% reduction in EPCR binding by the CIDRα1 domain. Antibodies purified by affinity for the CIDRα1 domain elicited a more than 90% reduction in EPCR binding at this concentration, while those purified by affinity for the synthetic binder caused an approximately 40% reduction. This suggests that sera raised through immunisation of rats with CIDRα1 domain contains antibodies which do not recognise the helix-kinked-helix motif and that these antibodies contribute significantly to prevention of EPCR binding.

We next conducted a similar experiment in which we purified human antibodies from a pool of polyclonal sera taken from fifteen semi-immune volunteers from a malaria endemic region of Tanzania, with sera selected based on their inhibitory effect on EPCR binding by HB3var03 CIDRα1.4 domain (Figure 3C). This antibody pool was tested for the ability to prevent EPCR binding by five different CIDRα1 domains, taken from five different domain subclasses. These antibodies, acquired in response to natural infection, had varying ability to prevent CIDRα1 domains from binding to EPCR. The binding of HB3var03 (CIDRα1.4) to EPCR was reduced by >90%, while the binding of IT4var20 (CIDRα1.1) and IT4var22 (CIDRα1.7) were reduced by ∼50%. A smaller inhibitory effect was seen for 1702_3 (CIDRα1.8) while there was no inhibitory effect on 1965_2 (CIDRα1.5).

We next affinity-purified antibodies from this human serum pool using a column coupled with the synthetic binder, and assessed the capacity of these antibodies to prevent CIDRα1 domains from binding to EPCR. In the case of IT4var20 CIDRα1.1 domain, the affinity purified antibodies retained about 40% of the inhibitory capacity of the original antibody mixture. In other cases, the majority of inhibitory antibodies were found in the run through of the column. These findings suggest that, in both CIDRα1 domain immunized rats, and in humans who have acquired antibodies as a result of natural exposure to parasites, the majority of antibodies that inhibit EPCR binding do not bind solely to the helix-kinked-helix motif. Nevertheless, antibodies which do bind to this region, found in the context of the synthetic binder, do block EPCR binding.

### The synthetic binder induces inhibitory, but not cross-reactive antibodies in immunized rats

As our synthetic binder is recognized by inhibitory antibodies from sera, we next assessed whether it would be effective as an immunogen to raise such antibodies. We immunized rats with the synthetic binder and assessed if the purified total antibodies from these sera block EPCR-binding by the HB3var03 CIDRα1.4 domain. Antibodies purified from rats immunised with synthetic binder caused a ∼40% decrease in EPCR binding by this CIDRα1 domain (Figure 4A). We again purified antibodies from this serum, either using a column coupled with the synthetic binder, or one coupled with the CIDRα1 domain. In this case, both columns depleted the majority of inhibitory antibodies from the serum, suggesting that the inhibitory antibodies present do target the helix-kinked-helix motif (Figure 4A).

**Figure 4:**
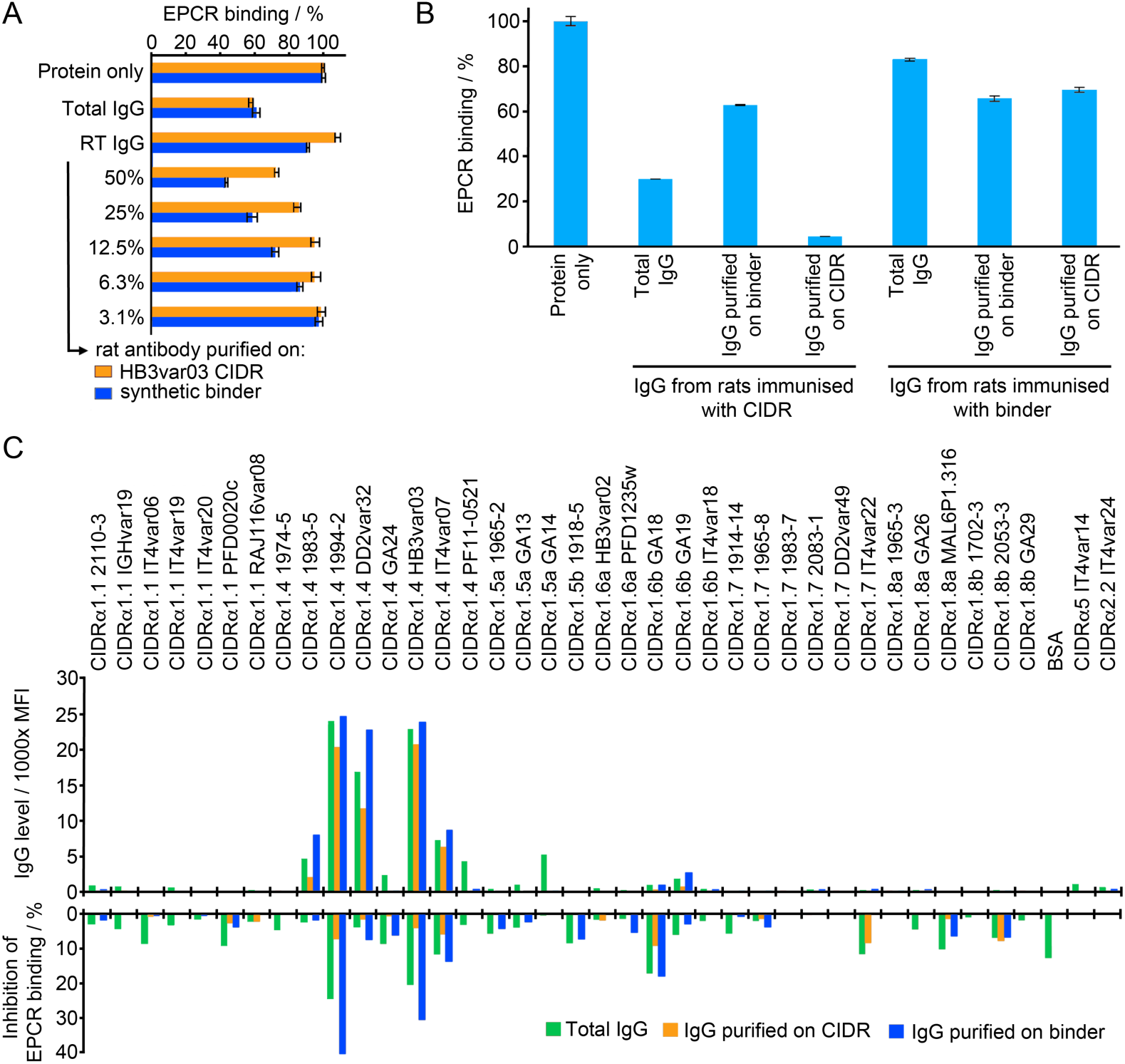
Immunogenicity of the synthetic binder. **A.** Antibodies from rats immunized with synthetic binder were affinity purified using either HB3var03 CIDRα1.4 domain or the synthetic binder. These were assessed for their ability to prevent CIDRα1.4 domain from binding to EPCR. Protein alone indicates binding of EPCR in the absence of antibody. Total IgG shows EPCR binding in the presence of 0.40mg/ml total IgG. RT IgG shows EPCR binding in the presence of antibodies that did not bind to the affinity column. The remaining columns show EPCR binding in the presence of affinity purified antibodies at different dilutions. All data are expressed as a percentage of the binding in the absence of antibody (protein only). **B.** Prevention of CIDRα1 binding to EPCR by rat antibodies was quantified at 25µg/ml. Antibodies were from rats immunized either with the synthetic binder or the HB3var03 CIDRα1.4 domain. Binding in the presence of Total IgG or of IgG affinity purified on the synthetic binder or the HB3var03 CIDRα1.4 domain was expressed as fraction of the binding in the absence of antibody (protein alone). **C.** IgG from rats immunized with synthetic binder was tested against a panel of CIDRα domains, either as total IgG, or after affinity purification on the synthetic binder or a HB3var03 CIDRα1.4 domain. The CIDRα2 and α5 domains are not expected to bind EPCR. The upper panel shows IgG binding levels (in mean fluorescence intensity, MFI). The lower panel shows the inhibition of the binding of these CIDRα domains to EPCR.

We next assessed, at an equivalent concentration of 25µg/ml, the effectiveness of these affinity purified antibodies when compared with those induced by immunization of a rat with CIDRα1 domain, and affinity purified on either CIDRα1 domain or synthetic binder (Figure 4B). This showed that, while immunization with the synthetic binder did generate antibodies which could inhibit CIDRα1 domains from binding to EPCR, these antibodies were less effective, both in pure serum, and at equal concentration, than those generated through immunization with CIDRα1 domains. This supports the view that antibodies which target sites other than the helix-kinked-helix motif are inhibitory or potentiate the effect of such inhibitory antibodies.

Finally, we assessed the breadth of reactivity of the antibodies generated through immunization with the synthetic binder. One of the goals of designing an immunogen containing the helix-kinked-helix motif was to attempt to raise more broadly reactive antibody mixtures, as this region of the domain is most conserved in chemistry and structure (8). We therefore tested the ability of the serum raised through immunization with the synthetic binder to recognize a panel of CIDRα1 domains from different subclasses in a Luminex assay (Figure 4C. Figure S6). In this case, we found that, as well as recognizing the cognate HB3var03 CIDRα1.4 domain, the sera recognized four other CIDRα1.4 domains. This cross-reactivity was retained in antibodies purified on either the synthetic binder or the HB3var03 CIDRα1 domain (Figure 4C). However, there was no reactivity to the other 31 CIDRα1 domains in the panel, or to the two CD36-binding CIDR domains. In parallel, we conducted a similar experiment in which we expressed the synthetic binder with a spy-tag at the C-terminus and coupled it to a spy-catcher conjugated virus like particle, before immunization and testing. However, this revealed no increase in breadth of reactivity (Figure S6). Therefore, antibodies induced through immunization with the synthetic binder were not more cross reactive than those generated through immunization with CIDRα1 domains (19).

## Discussion

A recent approach in vaccine design is to use structural methods to understand how critical protective monoclonal antibodies bind to a vaccine target and to then design a synthetic protein which recapitulates the structural features of the antibody epitope (22, 23). This has been used in the past to design novel vaccine components and to specifically re-elicit broadly inhibitory antibodies through immunization (24–27). In this study, we attempted a variant of this approach. In the absence of inhibitory antibodies against EPCR-binding PfEMP1, we instead used our knowledge of the structural features of the EPCR binding site (8), and grafted the core of this surface onto a three-helical bundle, which was redesigned through a Rosetta-based approach, to produce a small synthetic protein which mimics the key features of the EPCR binding surface. As the mimicked motif is the most conserved and functionally important surface feature of this protein family, we reasoned that antibodies that bind this region have the potential to be inhibitory and cross-reactive against the large and sequence diverse spectrum of EPCR-binding PfEMP1.

The design and production of the synthetic protein was successful. Surface plasmon resonance measurements showed that it bound to EPCR with a high nanomolar affinity matching that predicted. In addition, a crystal structure of the synthetic protein in complex with EPCR showed that it adopted a structure close to that expected, and bound in a mode that matched that of the parent CIDRα1 domain. We have therefore generated a synthetic protein which contains the core structural features of an EPCR-binding PfEMP1 domain.

We next tested the ability of this synthetic binder to interact with antibodies from sera from animals immunized with CIDRα1 domains, or from human volunteers from a malaria endemic region of Tanzania. We found that both sera contained antibodies which bound to the synthetic protein and that a fraction of these antibodies were inhibitory of EPCR binding. However, these antibodies were less effective than those purified through binding to the CIDRα1 domain, both in quantity and quality.

In addition, we immunized rats with either the HB3var03 CIDRα1 domain or the synthetic binder and assessed their capacity to recognize multiple CIDRα1 domains and to block their binding to EPCR. Immunisation of rats with the synthetic binder did generate antibodies which bind to their cognate CIDRα1 domain and prevent it from binding to EPCR. However, these were less effective and less abundant than equivalent antibodies raised through immunization with a CIDRα1 domain. In addition, when tested against a panel of CIDRα1 domains, the antibodies raised through immunization with the synthetic binder showed no more cross-reactivity than those raised through immunization with a CIDRα1 domain. Therefore, there is no evidence to suggest that focusing the immune response onto the helix-kinked-helix generates more cross-reactive and cross-inhibitory antibodies in rodents.

These findings suggest that many of the epitopes for inhibitory antibodies on the EPCR-binding CIDRα1 domains are not solely located on the helix-kinked-helix motif. This study therefore cautions against an approach in which a binding surface is grafted onto a smaller scaffold for immunization. In contrast, knowledge of the structure of the epitope of an effective antibody known to be elicited through human immunization is a valid starting point for such a study, allowing certainty that a complete epitope is recapitulated and also that it is possible to elicit such an antibody from the human germline. The quest to generate an epitope-focused vaccine targeting an EPCR-binding PfEMP1 protein therefore requires isolation and detailed molecular characterization of human monoclonal antibodies that target these domains as a starting point for future structure-guided vaccine development efforts.

### Experimental Procedures

#### Rosetta-based design

Epitope grafting used the Rosetta package (31), and was based on the fold-from-loops protocol (25). A composite model was generated in which the helix-kinked-helix motif was manually inserted into the three-helical bundle of PDB code 3LHP, chain S in Coot (29). An initial model was created by folding the helical bundle sequence around the fixed epitope using Cα-Cα restraints derived from the initial model. The resultant models were scored based on their Rosetta score and their root mean square deviation from the starting model. Sequence design was performed using a script from RosettaScripts (32), allowing all residues to change except for 72, 73, 75, 76, 77, 79, 80, 81, 82, 84 and 93. Four rounds of sequence design were conducted, using Cα-Cα restraints derived from the initial model. Ten thousand output models were generated and were filtered using Rosetta score. Those with scores lower than the starting model were then filtered using the packstat filter (33). As a final step, the best packed models were relaxed in the absence of Cα-Cα restraints and were filtered again using the packstat filter.

#### Protein Production

The HB3var03 CIDRα1.4 domain and its mutants were expressed in the BL21 strain of *E. coli* (8). The gene for the CIDRα domain was available in a modified pEt15b vector with an N-terminal hexa-histidine tag and a TEV cleavage site. Mutagenesis was performed using the Quikchange method to produce the D576A and K642A single and double mutants. Transformed *E. coli* were grown to an optical density of 1.0 at 600nm wavelength and expression was induced at 27°C by addition of IPTG to a final concentration of 1mM. Cells were harvested 3 hours after induction. The CIDRα1 domains expressed in the form of inclusion bodies, which were unfolded by incubation at room temperature in 6 M guanidine-hydrochloride, 20 mM Tris pH 8, 300 mM NaCl, 15 mM imidazole for 15 h. Refolding was achieved by gradual buffer exchange into 20 mM Tris pH 8, 300 mM NaCl, 15 mM imidazole in the presence of a glutathione redox buffer (3 mM reduced glutathione, 0.3 mM oxidised glutathione) while the protein was bound to a Ni-NTA column. Refolded protein was eluted and further purified by size exclusion gel chromatography (HiLoad Superdex 75 16/60, GE Healthcare) into 20 mM HEPES pH 7.5, 150 mM NaCl.

CIDRα1 domains used for immunization and EPCR binding assays, were produced in baculovirus-infected High Five cells as His_6_-tagged 30 kDa proteins (8) or 19 kDa STREP-II tagged proteins lacking the N-terminal β−sheet (19).

Designed synthetic binders were produced in *E. coli*. Synthetic genes were codon optimization for *E. coli* (GeneArt) and were inserted into the pEt15b vector to give an N-terminal His_6_ tag and a TEV cleavage site. Cysteine mutations to introduce disulphide bonds were incorporated using Quikchange mutagenesis (I10C L74C for Cys1; L53C L115C for Cys2 and V71C A100C for Cys3). The final synthetic binder was also cloned with a C-terminal spy-tag to allow attachment to virus-like particles decorated with spy-catcher (34, 35). The designed proteins were all expressed in inclusion bodies, by growth at 25 °C overnight after induction with 1mM IPTG. Protein was purified using the same on-column refolding method as for the CIDRα1 domains.

EPCR was expressed from a stable *Drosophila* s2 cell line, generating residues 16 to 210 fused to an N-terminal BAP tag, a His_6_-tag and a TEV cleavage site (8). Culture media was buffer exchanged into 20 mM Tris pH 8, 500 mM NaCl and protein was purified by Ni-NTA affinity chromatography and size exclusion gel chromatography (HiLoad Superdex 75 16/60, GE Healthcare) using 20 mM HEPES pH 7.5, 150 mM NaCl. Protein for crystallography was deglycosylated by treatment with endoglycosidase H_f_ (Sigma) and endoglycosidase F3 at enzyme:protein ratios of 1:50 in 50 mM MES pH 6.5 for 15 h. N-terminal tags were cleaved using TEV protease at an enzyme:protein ratio of 1:50 in PBS (Melford) with 3 mM reduced glutathione, 0.3 mM oxidised glutathione for 15 h at 25°C.

#### Surface plasmon resonance analysis

EPCR was biotinylated on the BAP tag by incubating 1 mg EPCR (30 μM) in 20 mM HEPES pH 7.5, 150 mM NaCl with 20 μg BirA, 0.3 μM biotin, 5 mM ATP, for 15 h at 25°C and was then coupled to CAPture chip (GE healthcare). This strategy was designed to allow EPCR to be immobilised with an orientation matching that found on the endothelial surface, and to generate a surface that could readily be regenerated.

SPR experiments were carried out on a Biacore T200 instrument (GE Healthcare). All experiments were performed in 20 mM HEPES pH 7.5, 150 mM NaCl, 0.005% Tween-20 at 25°C. Two-fold dilution series of each CIDRα domain or synthetic binder were prepared for injection over an EPCR-coated chip. For each cycle, biotinylated recombinant EPCR was immobilised on a CAP chip using the Biotin Capture Kit (GE Healthcare) to a total loading of 150 RU. Binding partners were injected for 240 s with a dissociation time of 300 s. The chip was regenerated in between cycles using regeneration solution from the Biotin Capture Kit (GE Healthcare). The specific binding response of the synthetic binders to EPCR was determined by subtracting the response given by the binder from a surface to which no EPCR had been coupled. The kinetic sensorgrams were globally fitted to a 1:1 interaction model to allow calculation of the association rate constant, k_a_; the dissociation rate constant, k_d_; and the dissociation constant K_D_ using BIAevaluation software version 1.0 (GE Healthcare).

#### Circular dichroism analysis

Circular dichroism measurements were taken using a J-815 spectrophotometer (JASCO) with attached Peltier water bath. Proteins were buffer exchanged into 10mM phosphate pH 7.5, 150mM NaF and were held in a quartz cuvette. Buffer-subtracted spectra were collected at wavelengths from 190nm to 260nm, at 25°C with ten spectra averaged for each measurement. For thermal melt experiments, spectra were taken at 2°C intervals from 20°C to 90°C.

#### Crystallisation and structure determination

For crystallization, TEV-cleaved synthetic binder was mixed with EPCR that had been TEV cleaved and deglycosylated at a molar ratio of 1.1 to 1, synthetic binder to EPCR. The complex was separated by size exclusion chromatography using a Superdex 200 16/60 column (GE Healthcare) into a buffer containing 20mM HEPES, 150mM NaCl pH 7.5 and was concentrated to 10.7mg/ml. Crystals grew in sitting drops with a well solution of 2M sodium citrate, 0.1M HEPES pH 7 at 4°C. Crystals were cryo-protected by transfer into 2M sodium citrate, 0.1M HEPES pH 7, 25% glycerol and were cryo-cooled in liquid nitrogen.

A complete data set was collected to 3.11Å resolution and data were indexed and scaled using Xia2 (36). Molecular replacement was performed in Phaser (37) using the structure of EPCR (PDB: 4V3D) (8) as a search model, identifying two copies in the asymmetric unit. Refinement in BUSTER (38), using the missing atoms functionality, revealed electron density corresponding to a single copy of the synthetic binder in complex with one of the copies of EPCR. A cycle of model building and refinement in COOT (39) and BUSTER allowed completion of the model of the synthetic binder, except for residues 35-50, which remained disordered.

#### Coupling to virus-like particles and immunization of rats

Synthetic binder was coupled to Acinetobacter bacteriophage AP205 virus like particles (VLPs) displaying one N-terminal SpyCatcher per capsid subunit (35). VLPs were expressed in *E. coli* BL21 StarTM (DE3) cells (Thermo Scientific) and purified by ultracentrifugation using an OptiprepTM density gradient (Sigma). Assembled VLPs and synthetic binder antigen were mixed at a 1:1 molar ratio and incubated for two hours at room temperature.

Assembled VLPs were quality assessed by Dynamic Light Scattering (DLS). The vaccine was centrifuged at 15,000 g for 10min, and the supernatant was loaded into a disposable Eppendorf Uvette cuvette (Sigma-Aldrich, USA) and measured 20 times at 25°C using a DynoPro NanoStar (WYATT Technology, USA) with a 658nm wavelength laser. Intensity-average size (nm) and percentage polydispersity (%Pd) were estimated using Dynamic software (Version 7.5.0).

Groups of four rats were immunized with 19 kDa HB3var03 CIDRα1.4 STRPII tagged protein (19). Groups of four rats were immunized with synthetic binder coupled to VLPs or synthetic binder alone. In all groups, rats received 20 µg synthetic binder. The rats were immunized intramuscularly (i.m.) every third week in a prime boost setting with Freund’s incomplete adjuvant for a total of three immunisations.

#### Immunoglobulin purification

One millilitre of serum was taken from each rat. From each immunization group of four rats, these were pooled and IgG was purified using protein G sepharose and was eluted into PBS to a total volume of 4 ml. 1750 µl of purified total IgG was passed over NHS columns loaded with 0.5 mg of either His_6_-tagged 30 kDa HB3var03 CIDRα1.4 or synthetic binder protein. Bound IgG was eluted and concentrated to 500 µl in PBS, and the inhibitory effect of these preparations were tested at different dilutions (Figure 3A, 4A and 4C). Alternatively, these purified IgG were further concentrated using Vivaspin columns before testing at equal protein concentrations (Figure 3B and 4B). Human IgG was similarly purified from plasma obtained from fifteen malaria-exposed Tanzanian donors selected by the ability of the plasma to inhibit HB3var03 CIDRα1.4 from binding to EPCR by ELISA.

#### Assessment of CIDRα domain reactivity and of the inhibition of EPCR-binding by CIDRα1 domains

An ELISA based assay was used to assess the inhibition of the binding of His_6_-tagged 30 kDa recombinant HB3var03 CIDRα1.4 (8) to EPCR. ELISA plates were coated with 3μg/mL recombinant EPCR, overnight at 4°C and were blocked with phosphate buffered saline (PBS) containing 3% skimmed milk. CIDRα1 protein was added at 5μg/ml concentration, with or without the addition of antibodies, and was incubated for one hour before washing three times with PBS containing 0.05% Tween-20. EPCR binding was determined using HRP conjugated anti-His_6_ antibody (1:3000). All ELISA assays were conducted to reach optical densities between 0.9 and 1.3 for the positive control without antibody addition and data is presented with this control normalized to 100%.

To assess cross-reactivity, IgG binding to a panel of 38 CIDRα domains coupled to Luminex microspheres, was measured (40). Serum was diluted 1:20, and IgG reactivity was detected using secondary phycoerythrin (PE)-conjugated antibody diluted to 1:3000. To assess inhibition of EPCR binding, microspheres were incubated with IgG at 1:50 for 30 minutes at room temperature. After washing with standard Luminex buffers, microspheres were incubated with 4 µg/mL biotinylated recombinant EPCR for 30 minutes at room temperature. EPCR binding was detected using PE-conjugated streptavidin.

## Data availability

Data for the structure reported here has been deposited in the PDB under the accession code 6SNY. Additional data supporting the findings reported in this manuscript are available from the corresponding author on request.

## Acknowledgments

The authors are grateful for the assistance of Ed Lowe and David Staunton (Department of Biochemistry, University of Oxford). We would also like to acknowledge the beamline staff at I02 for support during data collection. MKH was funded by Wellcome Trust project grant (087692/Z/08/Z). NMB was funded by a Wellcome funded Ph.D. studentship in Cellular Structural Biology. CKYL was funded by a Medical Research Council Ph.D. studentship. LT and TL were funded by Novo Nordisk Foundation (NNF16OC0023362, NNF17OC0029344, vNNF16OC0023056) and the Danish Council for Independent Research, Sapere Aude program DFF–4004-00624B.

---

**Figure S1:**
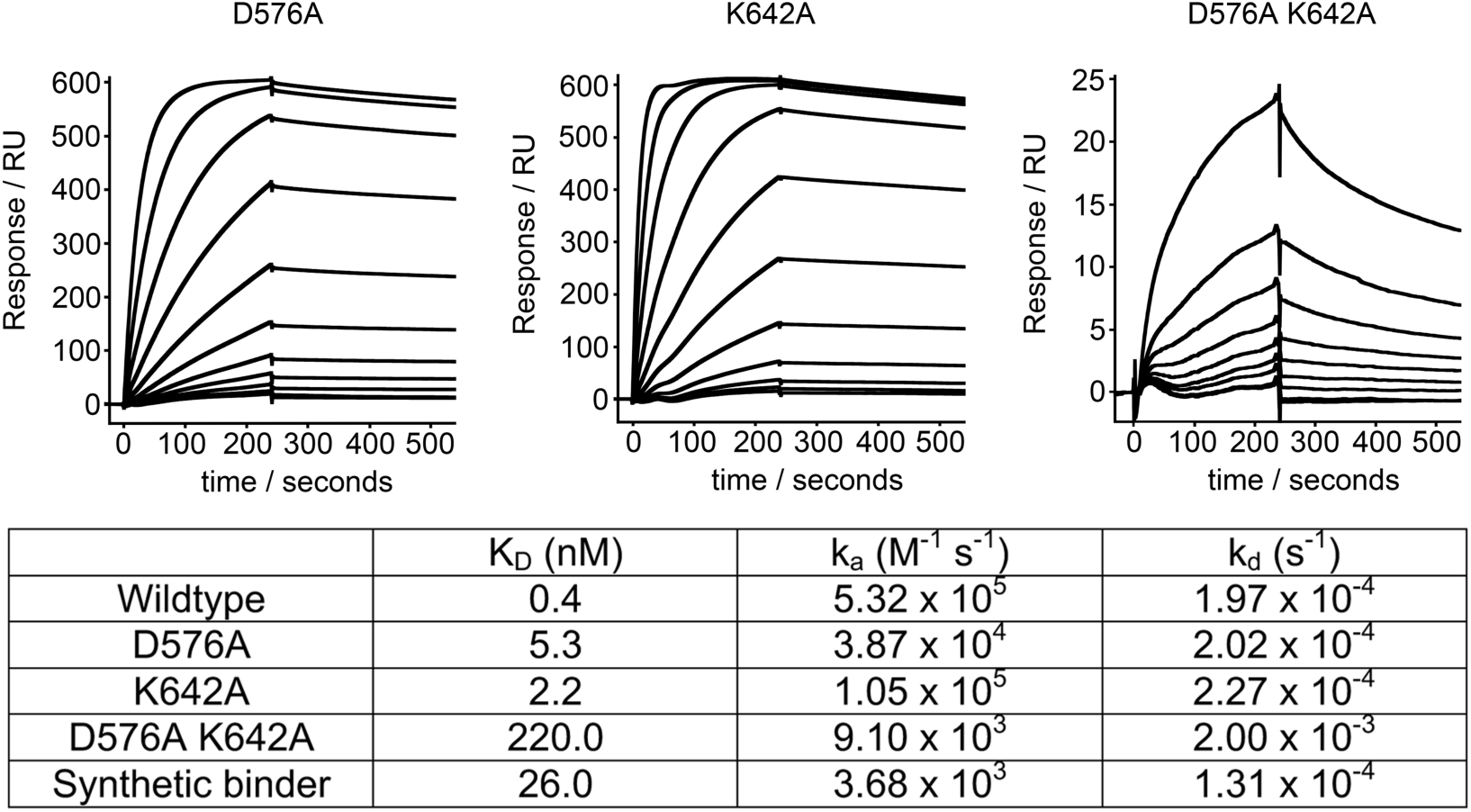
Residues on the helix-kinked helix are sufficient for EPCR binding. Surface plasmon resonance analysis of the binding of mutants of the HB3var03 CIDRα1 domain to EPCR. Nine residues from HB3var03 CIDRα1.4 make direct contacts to EPCR. Of these, only D576A and K642A are not present on the helix-kinked-helix. For both D576A and K642A, the curves show dilution series from 1μM to 0.9nM while for the D576A K642A double mutant, the curves show a dilution series from 250nM to 0.9nM. The table shows a comparison of the binding kinetics of the mutants to the wildtype domain, with the wildtype data from (8) and the parameters for the synthetic binder derived from the data presented in Figure 2A.

**Figure S2:**
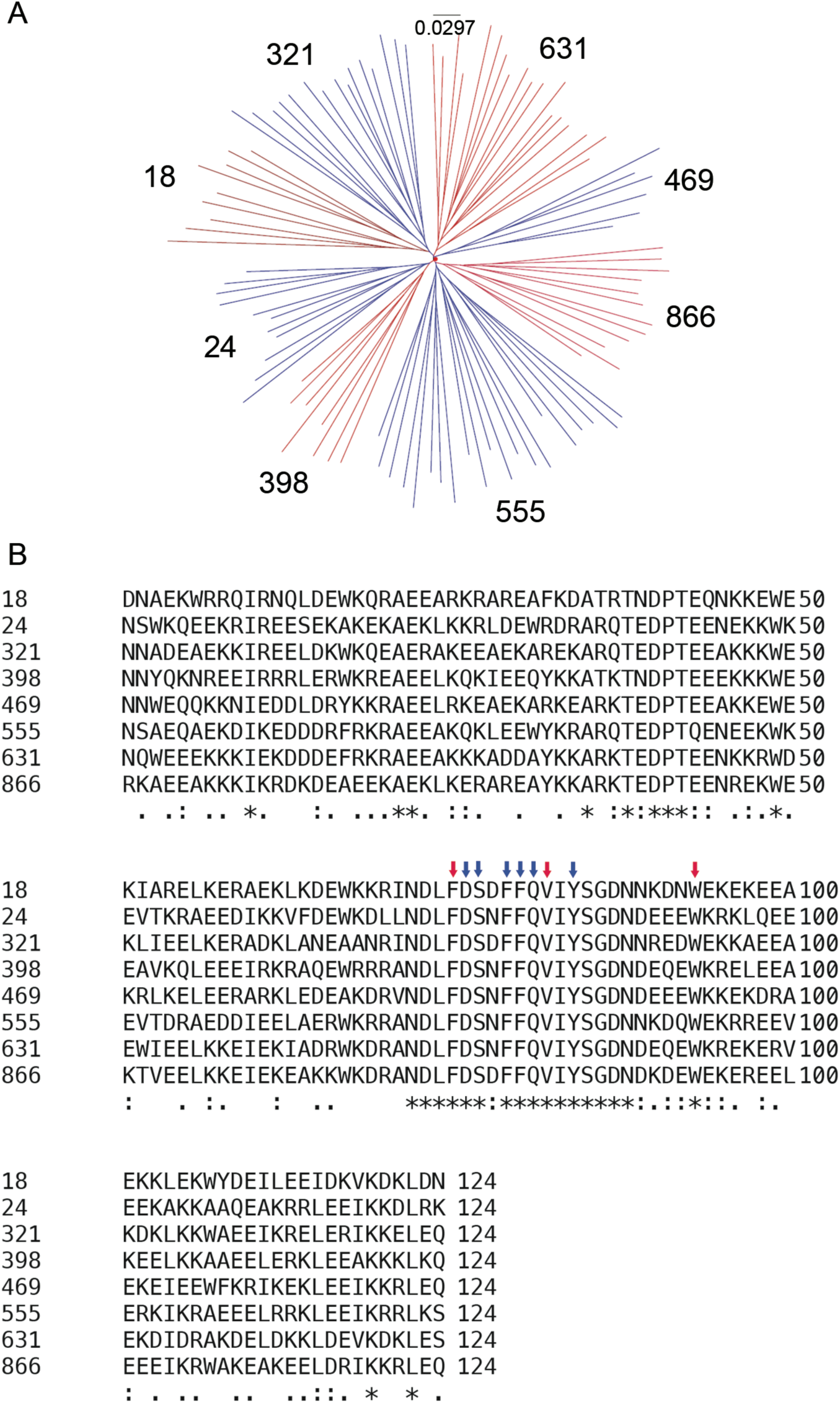
Selection of a panel of synthetic binder designs. **A.** A cladogram showing the diversity of the top 100 sequences. This was split into eight clades and one design was chosen from each clade. **B.** The sequences of the eight designs selected. The arrows represent residues which were fixed in sequence during the design process to match those in HB3var03 CIDRα1.4. The red arrows denote residues thought important in the formation of the kink in the kinked helix while the blue arrow indicates residues that contact EPCR directly.

**Figure S3:**
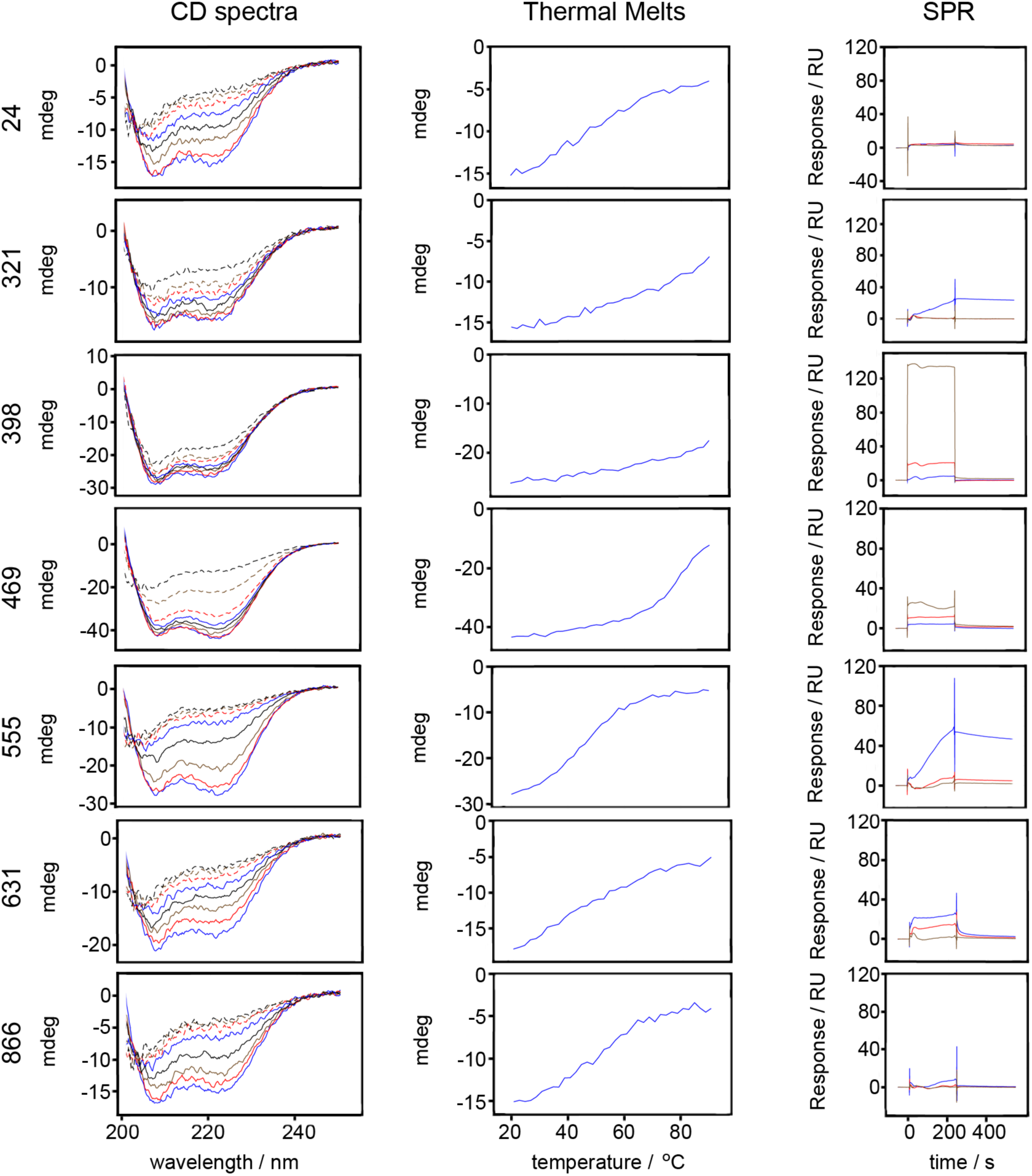
Analysis of the initial synthetic binder designs. The designs were tested for secondary structure content through their circular dichroism spectra (left), their thermal stability though the effect of temperature on their circular dichroism signal at 222nm (middle) and their binding to immobilized EPCR through surface plasmon resonance spectroscopy (right). The circular dichroism figure (left) shows data measured every 10°C from 20°C to 90°C. The surface plasmon resonance data (right) shows binding for 50μM (green), 5μM (red) and 0.5μM (blue) concentrations of each design to immobilized EPCR.

**Figure S4:**
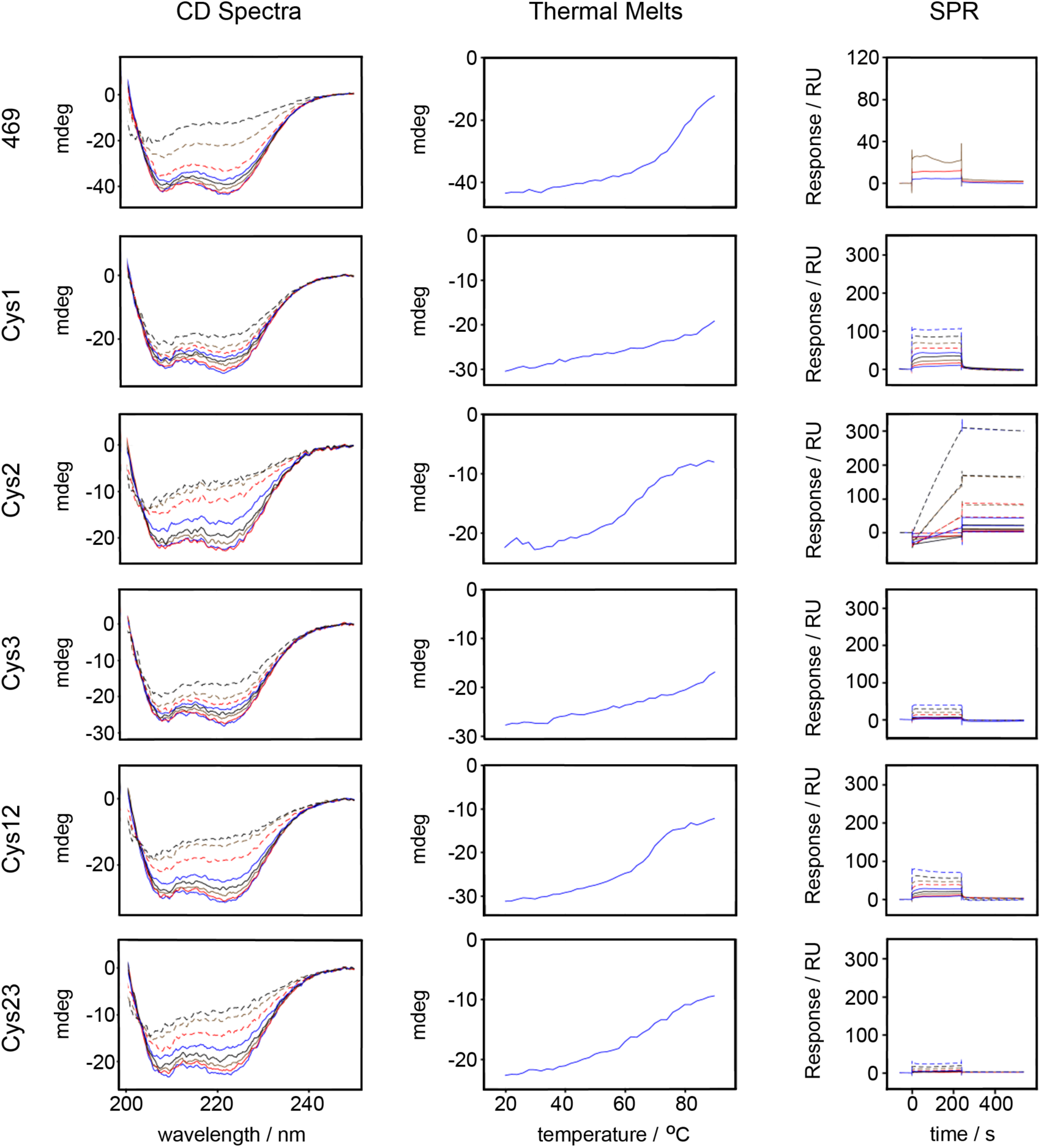
Analysis of the disulphide stabilized synthetic binders. Designs with added disulphide bonds were tested for secondary structure content through their circular dichroism spectra (left), their thermal stability based on the temperature dependence of their circular dichroism signal at 222nm (middle) and their binding to immobilized EPCR through surface plasmon resonance spectroscopy (right). The circular dichroism figure (left) shows the curves measured at every 10°C from 20°C to 90°C. The surface plasmon resonance data (right) shows binding for a two-fold dilution series of each design to immobilized EPCR, from a starting concentration of 12μM.

**Figure S5:**
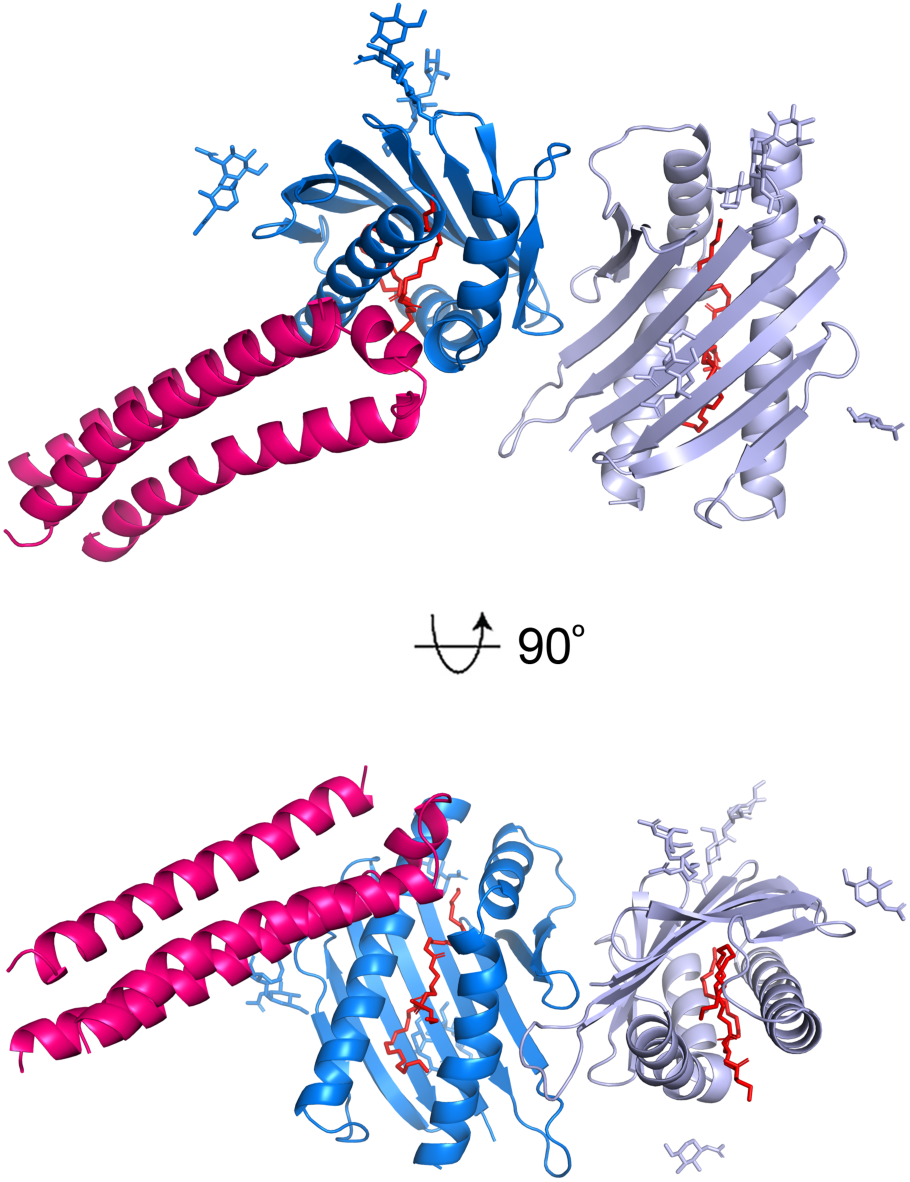
The asymmetric unit contains two EPCR and one synthetic binder. The synthetic binder is shown in pink, while the EPCR molecule interacting with the binder is blue. In light blue is shown a second EPCR which is not bound to a synthetic binder. Sugar molecules that form part of N-linked glycans are shown in sticks the same colour as the ribbons which represent the EPCR molecule to which they are attached. The lipids found within EPCR are represented as red sticks.

**Figure S6:**
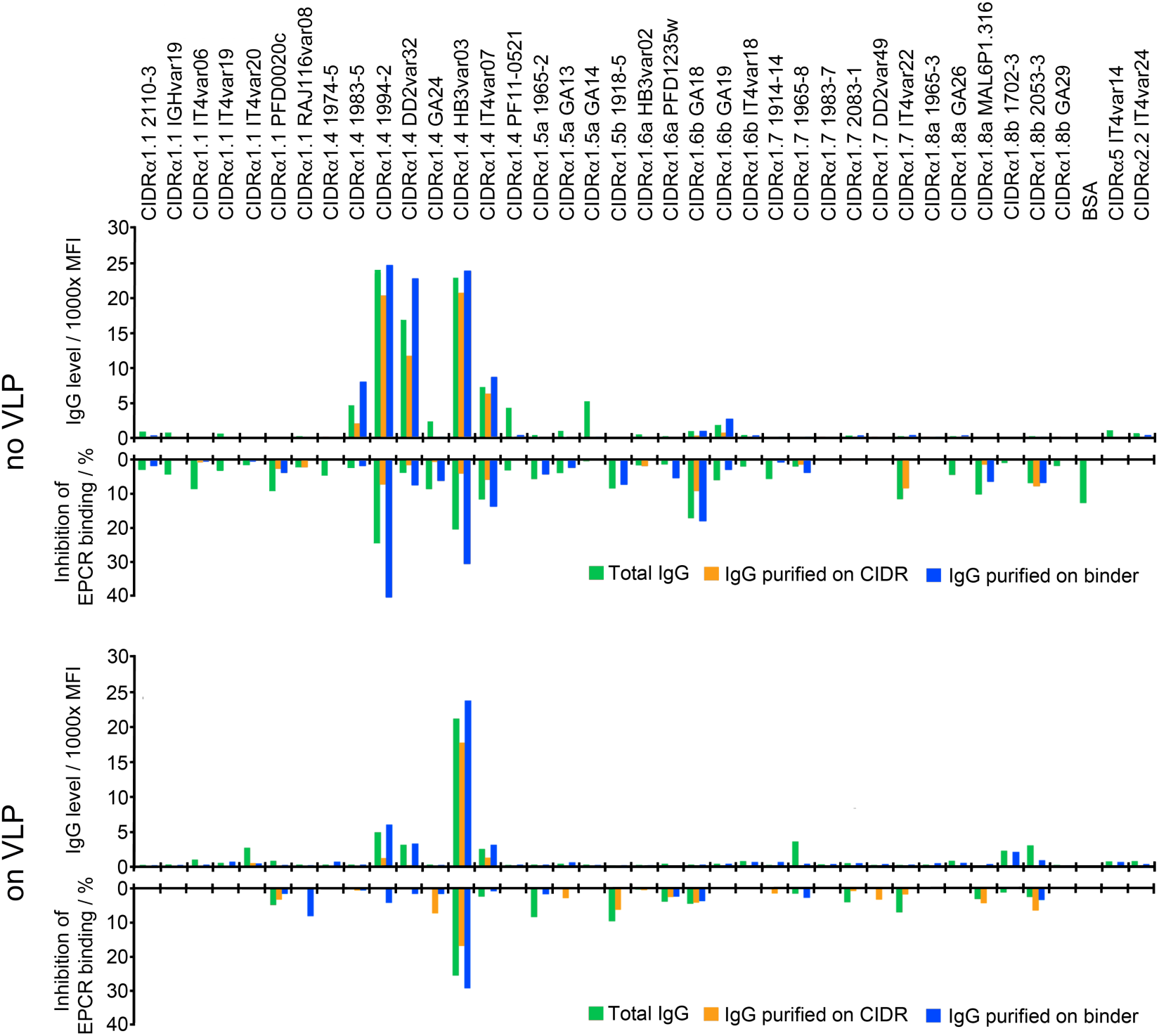
Comparison of the cross-reactivity of sera raised against the synthetic binding coupled and not coupled to virus-like particles. IgG from rats immunized with synthetic binder, either free (no VLP) or coupled to a virus-like particle (on VLP) was tested against a panel of CIDRα1 domains, either as total IgG (green bars), or after affinity purification on the HB3var03 CIDRα1.4 domain (yellow bars) or the synthetic binder (blue bars). The CIDRα2 and α5 domains are controls which do not bind EPCR. The upper panel shows IgG binding levels (in mean fluorescence intensity, MFI). The lower panel shows the inhibition of the binding of these CIDRα domains to EPCR.

**Table S1:**
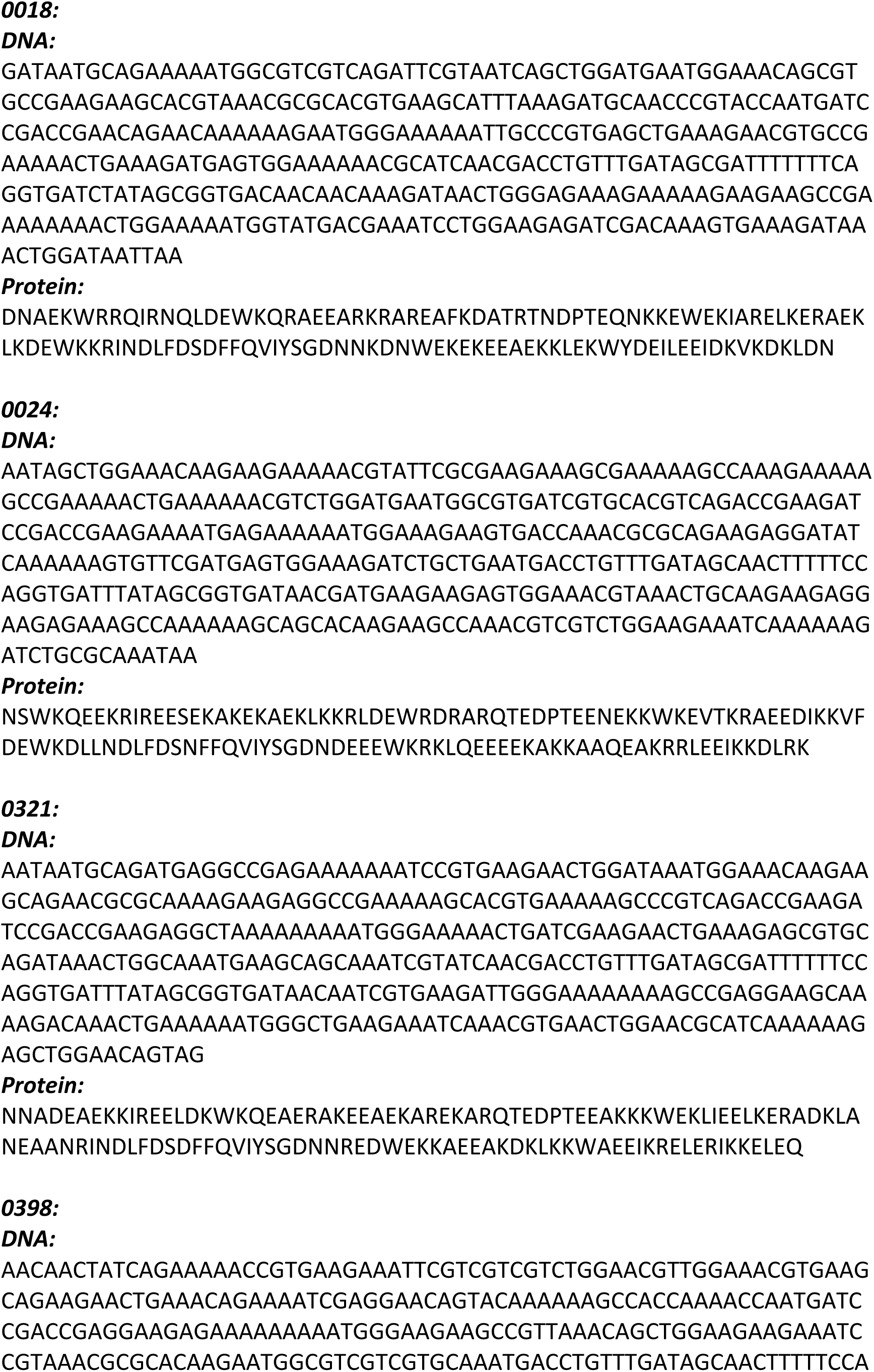
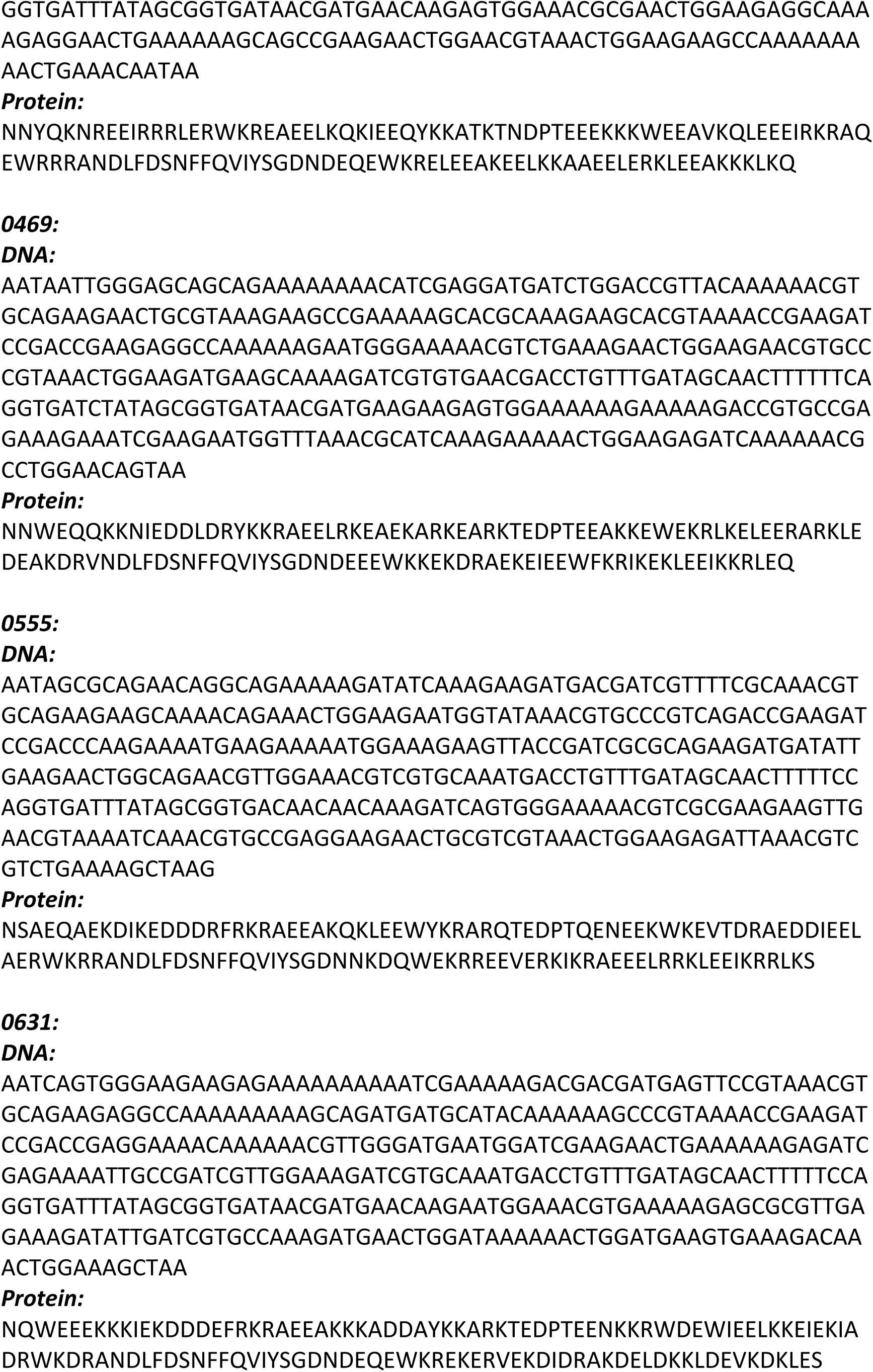
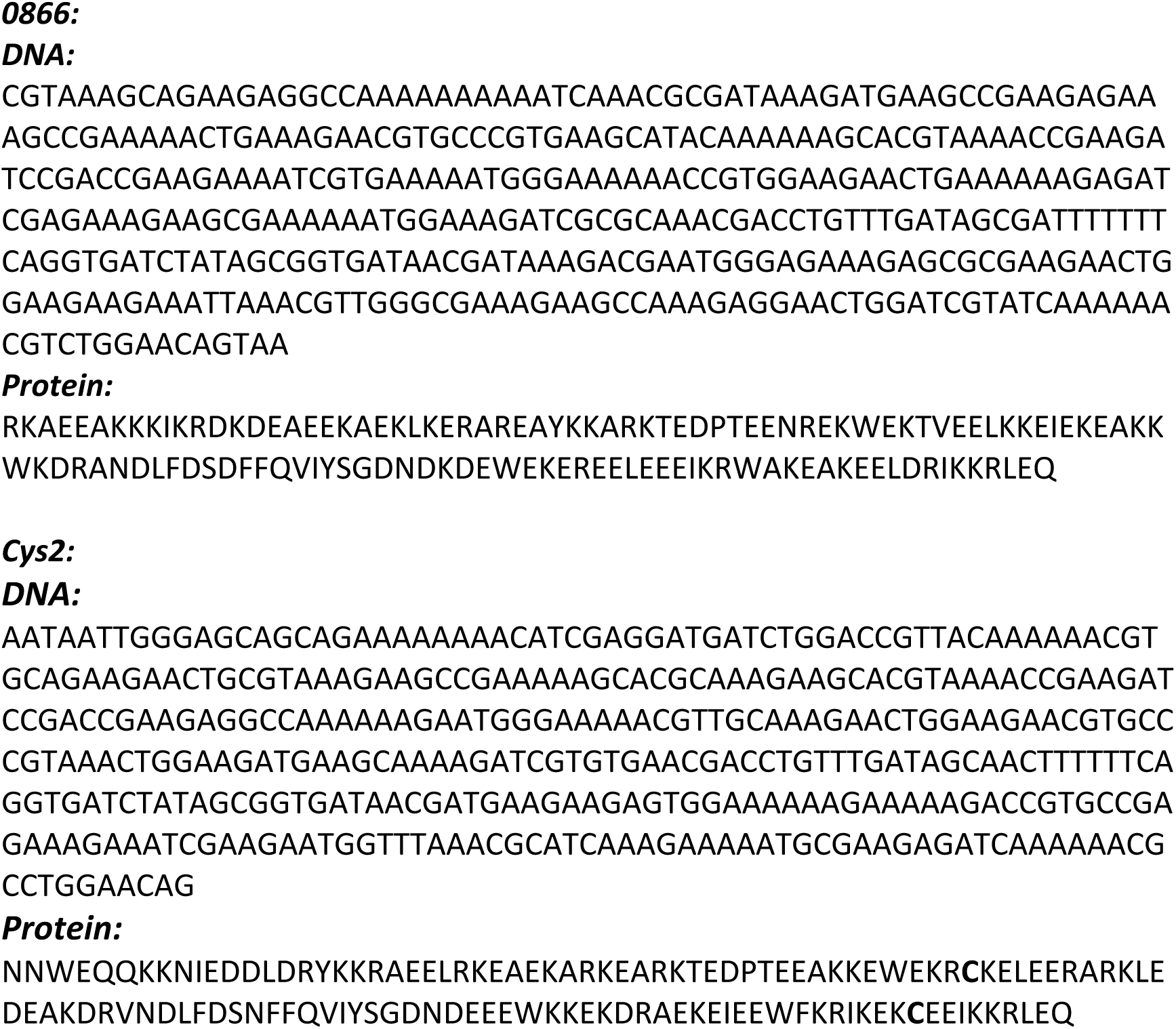
Sequences of synthetic binders:

**Table S2:**
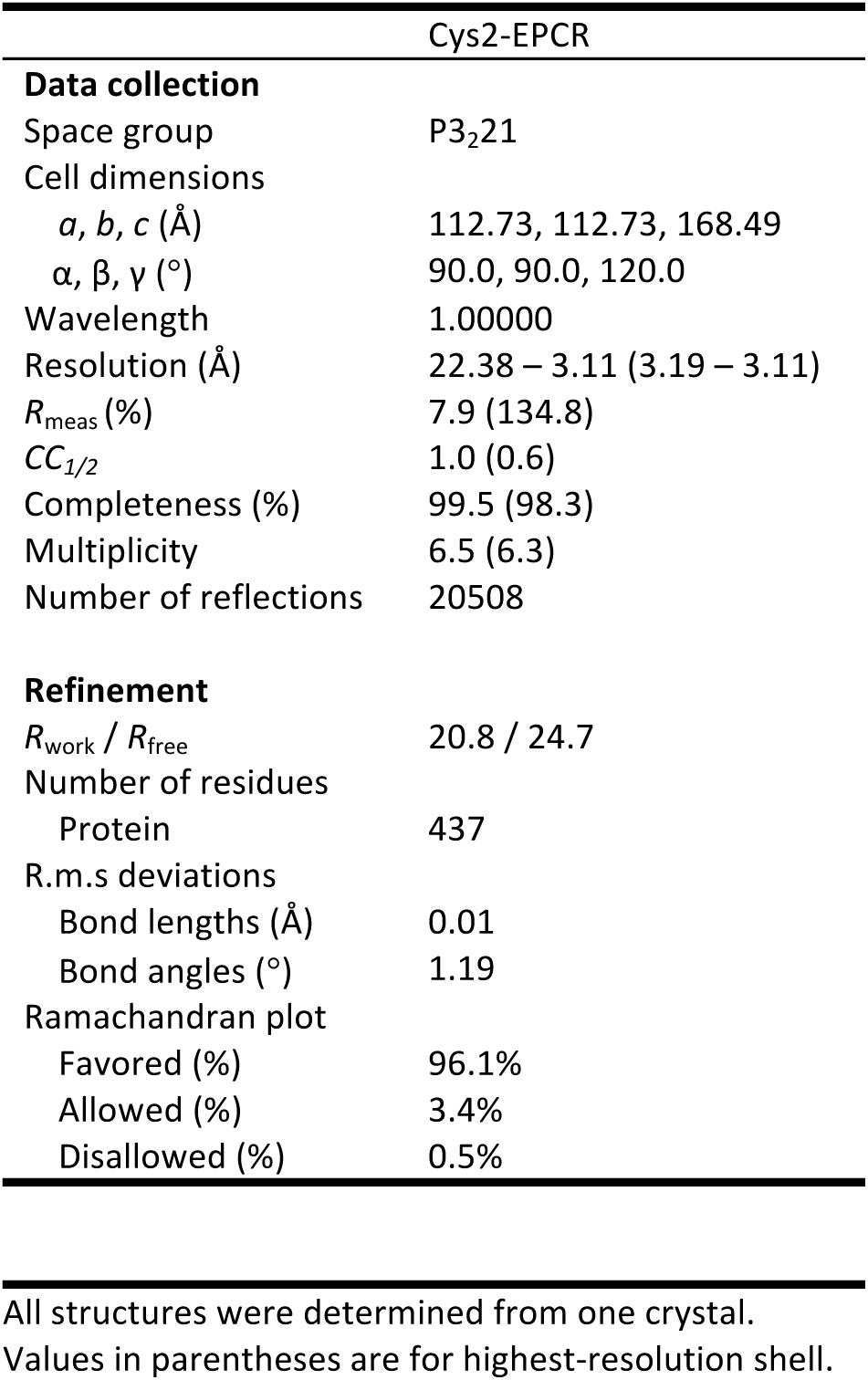
Data collection and refinement statistics.

